# Investigating the chemolithoautotrophic and formate metabolism of *Nitrospira moscoviensis* by constraint-based metabolic modeling and ^13^C-tracer analysis

**DOI:** 10.1101/2021.02.18.431926

**Authors:** Christopher E. Lawson, Aniela B. Mundinger, Hanna Koch, Tyler B. Jacobson, Coty A. Weathersby, Mike S. M. Jetten, Martin Pabst, Daniel Amador-Noguez, Daniel R. Noguera, Katherine McMahon, Sebastian Lücker

## Abstract

Nitrite-oxidizing bacteria belonging to the genus *Nitrospira* mediate a key step in nitrification and play important roles in the biogeochemical nitrogen cycle and wastewater treatment. While these organisms have recently been shown to exhibit metabolic flexibility beyond their chemolithoautotrophic lifestyle, including the use of simple organic compounds to fuel their energy metabolism, the metabolic networks controlling their autotrophic and mixotrophic growth remain poorly understood. Here, we reconstructed a genome-scale metabolic model for *Nitrospira moscoviensis* (iNmo686) and used constraint-based analysis to evaluate the metabolic networks controlling autotrophic and formatotrophic growth on nitrite and formate, respectively. Subsequently, proteomic analysis and ^13^C-tracer experiments with bicarbonate and formate coupled to metabolomic analysis were performed to experimentally validate model predictions. Our findings support that *N. moscoviensis* uses the reductive tricarboxylic acid cycle for CO_2_ fixation. We also show that *N. moscoviensis* can indirectly use formate as a carbon source by oxidizing it first to CO_2_ followed by reassimilation, rather than direct incorporation via the reductive glycine pathway. Our study offers the first measurements of *Nitrospira’s in vivo* central carbon metabolism and provides a quantitative tool that can be used for understanding and predicting their metabolic processes.

**Importance:** *Nitrospira* are globally abundant nitrifying bacteria in soil and aquatic ecosystems and wastewater treatment plants, where they control the oxidation of nitrite to nitrate. Despite their critical contribution to nitrogen cycling across diverse environments, detailed understanding of their metabolic network and prediction of their function under different environmental conditions remains a major challenge. Here, we provide the first constraint-based metabolic model of *N. moscoviensis* representing the ubiquitous *Nitrospira* lineage II and subsequently validate this model using proteomics and ^13^C-tracers combined with intracellular metabolomic analysis. The resulting genome-scale model will serve as a knowledge base of *Nitrospira* metabolism and lays the foundation for quantitative systems biology studies of these globally important nitrite-oxidizing bacteria.

## Introduction

The oxidation of nitrite to nitrate is a key step in nitrification and the global nitrogen cycle. The process is a critical control point counteracting nitrogen loss to the atmosphere and is mediated by a phylogenetically diverse functional guild known as the nitrite-oxidizing bacteria (NOB) (1). The genus *Nitrospira* constitutes the most diverse and abundant NOB based on marker gene (16S ribosomal RNA and nitrite oxidoreductase (NXR)) and metagenomic surveys. The genus *Nitrospira* consists of at least six lineages that mediate nitrite oxidation across various habitats, including soil, freshwater, marine, terrestrial and engineered ecosystems (1)(2). *Nitrospira* must be flexible enough to survive in the wide range of fluctuating environmental conditions characteristic of these habitats, suggesting their ecophysiology and ecological niches extend beyond those initially defined by their chemolithoautotrophic lifestyle.

Despite their recalcitrance to cultivation, recent insights driven by metagenomics have shed light on the unique features of *Nitrospira’s* carbon and energy metabolism (3). *Nitrospira* encode novel respiratory chain enzymes for energy conservation, including an evolutionarily distinct membrane-bound periplasmic nitrite oxidoreductase (NXR) and a putative cytochrome *bd*-like oxidase that may allow them to adapt to low dissolved oxygen environments (3). *Nitrospira* also encode all genes for CO_2_ fixation via the reductive tricarboxylic acid cycle (rTCA) and lack the two key genes (ribulose bisphosphate carboxylase and phosphoribulokinase) needed to operate the Calvin-Benson-Bassham cycle (CBB) common in other NOB (3). Outside their chemolithoautotrophic growth, genomic and experimental data have revealed that *Nitrospira* can use alternative substrates to fuel their carbon and energy metabolism (1)(4)(5). In addition to nitrite, some *Nitrospira* species have been experimentally shown to use formate, hydrogen, and ammonia as electron donors with oxygen or nitrate as terminal electron acceptors (4)(5)(6)(7)(8). For example, *N. moscoviensis* contains a soluble formate dehydrogenase and NiFe hydrogenase that allows growth with formate or H_2_, respectively (4)(5), whereas *N. inopinata* contains pathways for both ammonia and nitrite oxidation that enable growth via complete nitrification (8). In addition to their energy metabolism, fluorescence *in situ* hybridization combined with microautoradiography (FISH-MAR) experiments have also suggested that *Nitrospira* populations present within activated sludge microbial communities can assimilate pyruvate (2) and formate (9), although the carbon assimilation pathways for these substrates have yet to be determined. While this expanded metabolic versatility suggests that *Nitrospira* are adapted to dynamic environmental conditions, our ability to predict their function in natural and engineered ecosystems is constrained by the limited understanding of their metabolic network and the lack of quantitative tools to study their metabolism.

Genome-scale metabolic modeling is a powerful method for analyzing and predicting the biochemical pathways driving microbial metabolism. Such modeling approaches calculate the flow of metabolites through a reconstructed metabolic network based on relevant constraints (e.g. network stoichiometry, thermodynamics, measured fluxes) using a technique called flux balance analysis (FBA) (10). This provides a quantitative framework for analyzing metabolism and predicting phenotypes when combined with physiological data. Moreover, genome-scale models can be used to generate testable hypotheses on the functional capabilities of organisms under defined conditions that can subsequently be tested; for example, using ^13^C isotope tracing combined with metabolomics (11).

Here, we provide the first constraint-based metabolic reconstruction and analysis of *N. moscoviensis* representing the ubiquitous *Nitrospira* lineage II. We examine the metabolism of *N. moscoviensis* growing under canonical chemolithoautotrophic conditions and also during growth with formate as a substrate. Taking advantage of recent advances to cultivate *Nitrospira* in continuous flow membrane bioreactors (12), we subsequently validate *N. moscoviensis’* predicted metabolic network using proteomics and ^13^C-tracers combined with quantitative metabolomic analysis. Our proteomic and ^13^C metabolomic results support the use of the rTCA for carbon fixation by *N. moscoviensis* during chemolithoautotrophic growth. We further show that *N. moscoviensis* does not assimilate formate directly, but instead re-assimilates CO_2_ produced via formate oxidation using the rTCA cycle. The resulting genome-scale model (GEM), *iNmo686*, will serve as a knowledge base for understanding and predicting the function of *Nitrospira* in both natural and engineered ecosystems.

## Results

### Genome-scale metabolic reconstruction of Nitrospira moscoviensis

The genome-scale metabolic network of *N. moscoviensis* (*iNmo686)* was reconstructed from the most recent *N. moscoviensis* genome annotation (NCBI accession number NZ_CP011801), aided by reaction annotations in the MetaCyc (13) and ModelSEED databases available through the Department of Energy Systems Biology Knowledgebase (14). Proteomic analysis of *N. moscoviensis* grown on nitrite was also performed to confirm that model reactions carrying flux were detected in the proteome (Supplementary Table 1). The final reconstruction contained a total of 678 reactions, 638 metabolites, and 686 genes (Supplementary Dataset 1). Reactions for *Nitrospira’s* respiratory chain were reconstructed based on existing models of electron flow (1)(3) and assuming that the two protons produced during nitrite oxidation in the periplasm contribute to proton motive force generation. Cytoplasmic reactions for formate dehydrogenase and hydrogenase that catalyze formate oxidation to CO_2_ and hydrogen oxidation, respectively, were also included based on genomic and experimental evidence (4)(5). Formate transport and assimilation reactions mediated by a putative reductive glycine pathway (15) encoded in the genome (Supplementary Dataset 1) were also included to evaluate the possibility of directly assimilating formate as a carbon source, as suggested by previous FISH-MAR observations (9).

Since the stoichiometry for proton translocation in *Nitrospira’s* respiratory chain complexes is unknown, values were assumed from model organisms (Supplementary Dataset 1; Figure 1). The mechanism for reducing low-potential electron carriers, such as ferredoxin, required for carbon fixation in *Nitrospira* is also unknown, but it has been hypothesized that the 2M-type NADH dehydrogenase complex in *Nitrospira* performs this function (16). Thus, the model included this mechanism for ferredoxin reduction. All annotated biosynthetic and biodegradation reactions for amino acids, nucleic acids, carbohydrates, lipids, and cofactors were included in the reconstruction (Supplementary Datasets 1 & 2). This encompassed all predicted central carbon metabolic reactions annotated in the genome, including reactions of the rTCA cycle, gluconeogenesis, the pentose phosphate pathway, anaplerotic reactions, one-carbon metabolism, and fatty acid metabolism. Gaps in the network were identified and filled manually to ensure that *iNmo686* could grow on minimal NOB media (see Materials and Methods).

**Figure 1.**
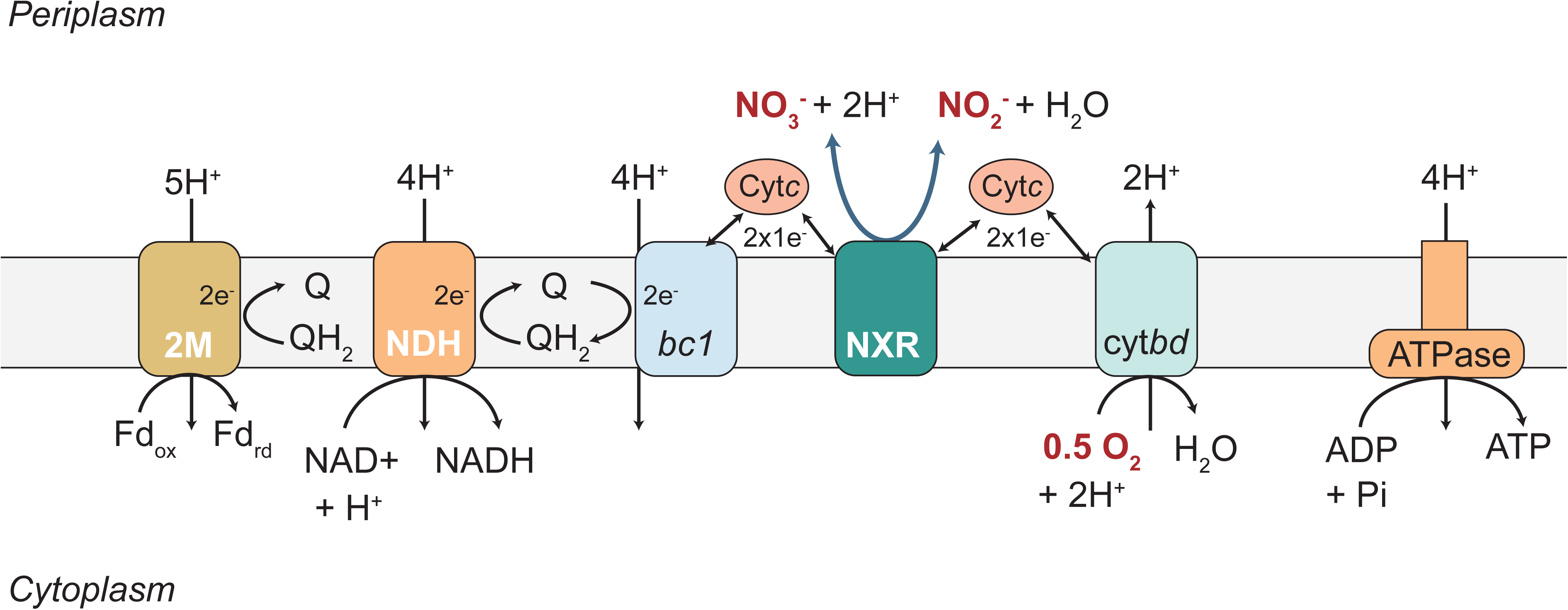
Theoretical model for the energy metabolism of *Nitrospira moscoviensis* hypothesized in this study. Q: quinone; QH2: quinonol; Fd: ferredoxin; 2M: 2M-type complex I; NDH: NADH dehydrogenase (complex I); bc1: cytochrome bc1 complex (complex III); NXR: nitrite oxidoreductase; cyt*bd*: cytochrome *c* oxidase (complex IV).

To obtain qualitative and quantitative outputs from a genome-scale model via FBA, an objective function is required. This is typically accomplished based on a biomass objective function, which assumes that maximization of biomass growth is the cellular objective (17). Therefore, to generate a representative biomass objective function for *N. moscoviensis* we experimentally determined its biomass composition (i.e. macromolecular components) during steady-state growth (see Materials and Methods).

A summary of the biomass composition for *N. moscoviensis* is presented in Tables 1 and 2. These measurements, together with the genome sequence data and published fatty acid composition data (18), were used to formulate *N. moscoviensis’* biomass objective function (Supplementary Dataset 1). Growth and non-growth associated maintenance energy requirements were estimated to be 535 mmol ATP gDW^-1^ and 0.90 mmol ATP gDW^-1^ hr^-1^, respectively, by plotting the experimentally determined nitrite uptake rate against the growth rate and using a net ATP yield of 1.0 mmol ATP mmol NO_2_-N^-1^ determined from the model (Supplementary Figure 1).

**Table 1.** N. moscoviensis biomass composition.

| Molecule | Average | Std Dev |
| --- | --- | --- |
| RNA | 2.4 | 0.04 |
| DNA | 1 | n.d. |
| Proteins | 46.7 | 4.7 |
| Cabohydra | 26.9 | 1.8 |
| Lipids | 16.3 | 3.2 |
| Inorganics | 5 | n.d. |

**Table 2.**
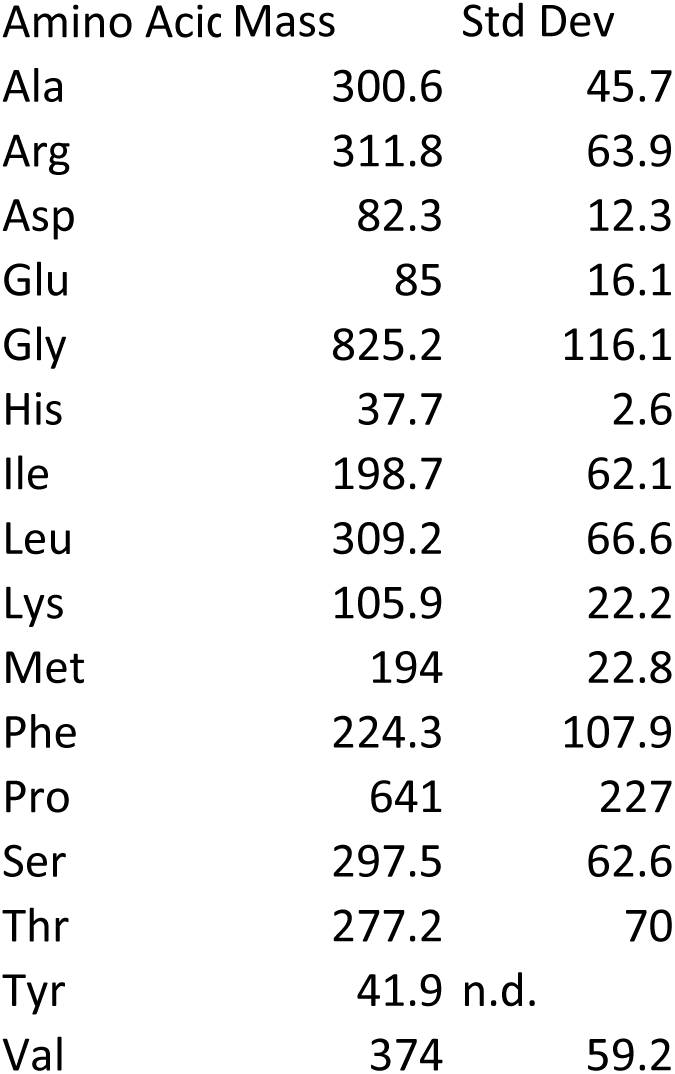
N. moscoviensis amino acid composition.

| Amino Acid | Mass | Std Dev |
| --- | --- | --- |
| Ala | 300.6 | 45.7 |
| Arg | 311.8 | 63.9 |
| Asp | 82.3 | 12.3 |
| Glu | 85 | 16.1 |
| Gly | 825.2 | 116.1 |
| His | 37.7 | 2.6 |
| Ile | 198.7 | 62.1 |
| Leu | 309.2 | 66.6 |
| Lys | 105.9 | 22.2 |
| Met | 194 | 22.8 |
| Phe | 224.3 | 107.9 |
| Pro | 641 | 227 |
| Ser | 297.5 | 62.6 |
| Thr | 277.2 | 70 |
| Tyr | 41.9 | n.d. |
| Val | 374 | 59.2 |

### Proteome of N. moscoviensis grown on nitrite

A high coverage proteome of *N. moscoviensis* grown on nitrite was obtained to confirm expression of the reconstructed metabolic network. Enzymatic digests using three different digestion enzymes alone and in combinations were tested to ensure a representative membrane proteome was obtained (see Supplementary Information for details). Whole cell proteome analysis resulted in the detection of 2519 of the 4733 non-identical proteins (53.2%) encoded in the *N. moscoviensis* genome, including 344 of the 878 non-identical membrane proteins (39.2%) (Figure 2; Supplementary Table 1). The resulting proteome confirmed expression of genes involved in *N. moscoviensis* carbon and energy metabolism represented in the model, including the nitrite oxidation system (NXR), the reductive TCA cycle, and nitrogen assimilation pathways (Supplementary Table 2).

**Figure 2.**
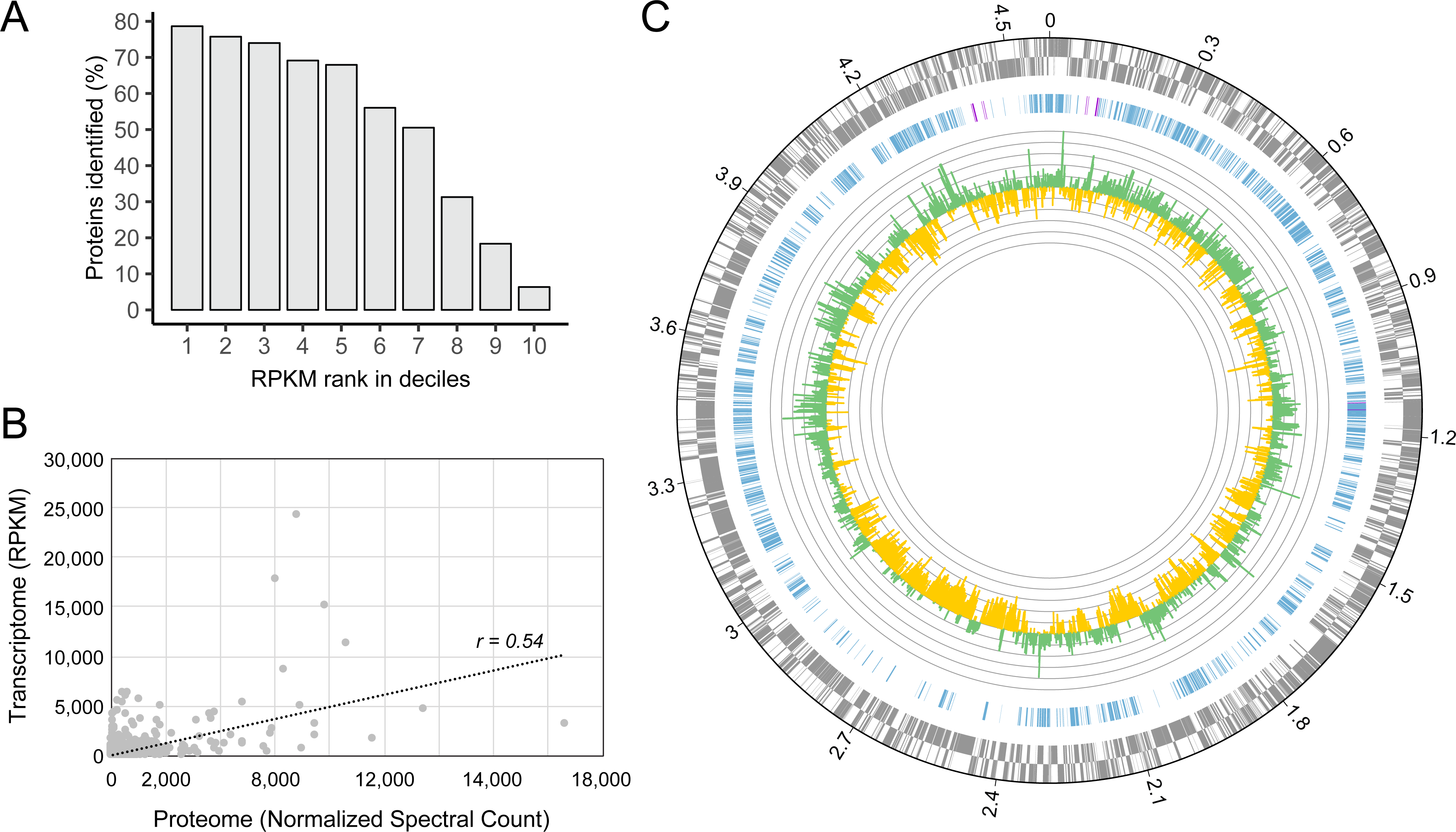
*Nitrospira moscoviensis* proteomic analysis. A) Number of proteins identified in the whole cell proteome given per unique uniprot identifier by their gene expression in deciles, ranked by RPKM from high (1) to low (10). B) Pearson correlation (*r*=0.54, *n*=2788) between open reading frame (ORF) transcript abundance (RPKM) and protein abundance (normalized spectral counts) across all ORFs with detected peptides (see Supplementary Table 1). C) Genome-wide proteomic and transcriptomic profile of N. moscoviensis during growth on nitrite. Rings are counted from outside to inside (1–3). (1) Genes (+/- strand) in the published N. moscoviensis genome (CP011801). (2) CDS corresponding to proteins identified in the whole cell proteome, CDS with 100% identity are marked in purple. (3) Average gene expression levels (n=3, with 3 technical replicates each) in log_2_-fold to median, with green representing transcription level above and yellow below the median shown in log_2_ values.

The expressed proteome qualitatively corresponded to the recently published transcriptome of *N. moscoviensis* (12), where 86% of all detected proteins also displayed gene expression levels above median (Figure 2; Supplementary Table 1). However, protein and transcript abundances were only moderately correlated when quantitatively compared using normalized spectral counts as a proxy for protein abundance (*r=0.54*; Figure 2B). While some of the highly transcribed coding sequences (CDS) that lacked detection on the proteome level represent erroneous gene callings (such as NITMOv2_0031, NITMOv2_0852), there were also proteins which, despite relatively high transcript levels, were not detected at all in the proteome, such as ATP-dependent RNA helicase RhlE (NITMOv2_1402) or the periplasmic phosphate-binding protein (PstS) of the phosphate ABC transporter (NITMOv2_4757). Such discrepancies can be caused by posttranscriptional regulation or limitations of the proteomic method, such as insufficient solubilization or the presence of few potential cleavage sides for the tryptic digest.

### Analysis of chemolithoautotrophic growth on nitrite

The *iNmo686* genome-scale model was first used to quantify the carbon and electron flux distribution in *N. moscoviensis* during aerobic growth on mineral media with nitrite as an electron donor and CO_2_ as a carbon source. The model was constrained by the nitrite uptake rate measured during chemostat cultivation, and the objective function was maximizing growth. Consistent with experimental data, when the nitrite uptake rate was set to 8.5 mmol NO_2_^-^ gDW^-1^ hr^-1^ the specific biomass growth rate equaled 0.006 hr^-1^ (∼116 hr doubling time; Supplementary Figure 1). Under these conditions, approximately 66% of the oxidized NO_2_^-^ is used for growth associated maintenance (GAM; polymerization of amino acids into proteins, RNA error checking, etc.) and 17% is oxidized for non-growth associated maintenance (NGAM; maintenance of chemical gradients, turgor pressure, etc.) in the form of ATP (Supplementary Figure 2). The remaining 17% is used to generate ATP (7%) and reducing equivalents (10%) required for CO_2_ fixation and synthesis of macromolecular building blocks (e.g. central metabolites, amino acids, nucleotides, fatty acids). Electron transport chain (ETC) reactions involved in nitrite oxidation carried the highest flux and were an order of magnitude higher than carbon fixation fluxes (Figure 3 and Figure 4). This agrees with *N. moscoviensis*’ large maintenance energy demand and the high abundance of ETC proteins measured in the proteome (Supplementary Table 2).

**Figure 3.**
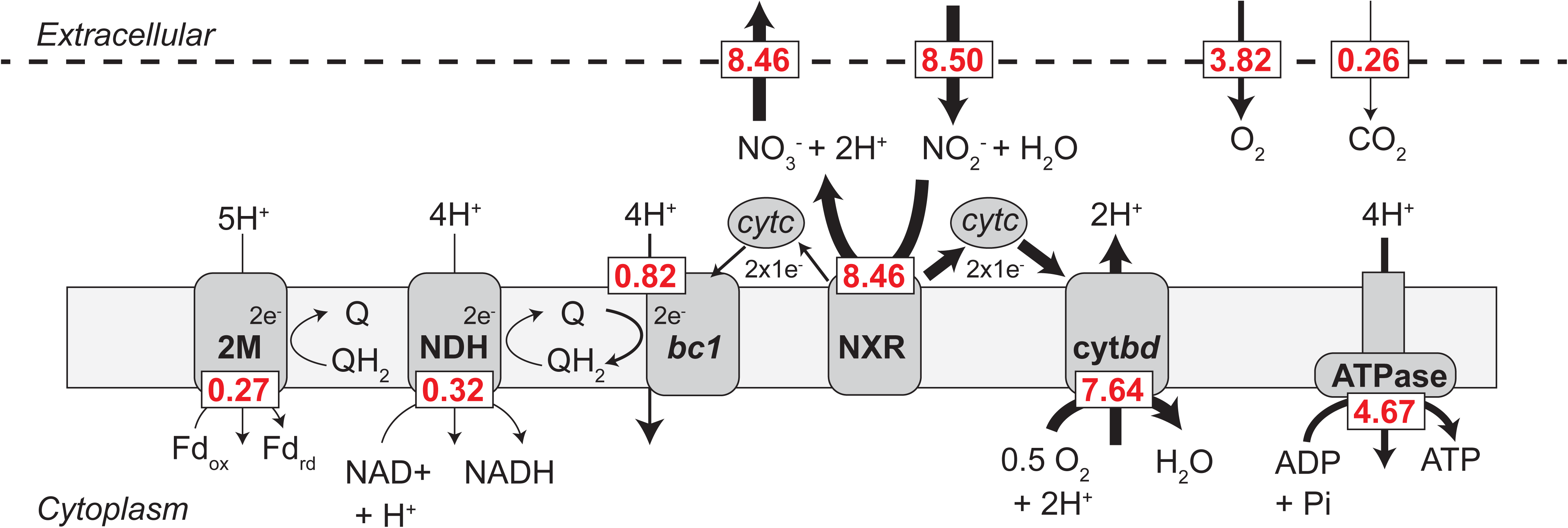
Electron flux distribution predicted via flux balance analysis during chemolithotrophic growth on nitrite. The model was constrained to a nitrite uptake rate of 8.5 mmol gDW^-1^ hr^-1^ and the biomass growth rate was 0.006 hr^-1^. Numerical values (red) are calculated fluxes in units of mmol gDW^-1^ hr^-1^. Model reactions and compounds can be found in Supplementary Dataset 1. FBA solutions can be found in Supplementary Dataset 2.

**Figure 4.**
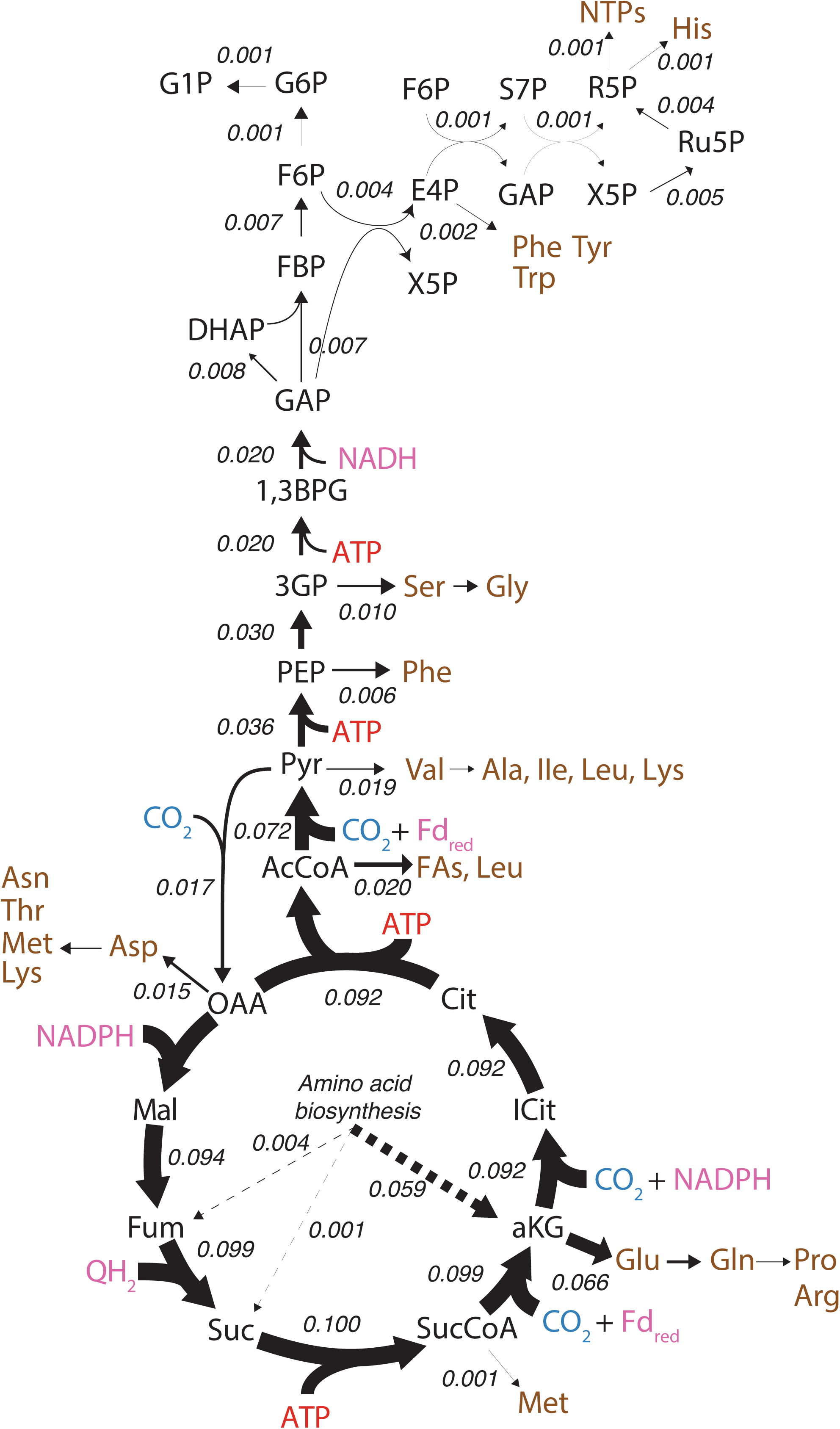
Carbon flux distribution predicted via flux balance analysis during chemolithoautotrophic growth on nitrite. The model was constrained to a nitrite uptake rate of 8.5 mmol gDW^-1^ hr^-1^ and the biomass growth rate was 0.006 hr^-1^. CO_2_ is shown in blue, amino acids and other biomass precursors are shown in brown, ATP is shown in red, and reducing equivalents are shown in pink. Numerical values are calculated fluxes in units of mmol gDW^-1^ hr^-1^. Model reactions and compounds can be found in Supplementary Dataset 1. FBA solutions can be found in Supplementary Dataset 2.

The predicted carbon flux distribution in *N. moscoviensis* during chemolithoautotrophic growth showed that enzymes of the rTCA cycle, including 2-oxoglutarate:ferredoxin oxidoreductase (OFOR), succinyl-CoA synthetase (SCS), ATP-citrate lyase, (ACL), and pyruvate:ferredoxin oxidoreductase (PFOR) carried the highest carbon flux, consistent with their primary role in CO_2_ fixation (Figure 4). Pyruvate carboxylase also carried high carbon flux (Figure 4), which is required to replenish TCA cycle intermediates used as precursors for biosynthesis (e.g. oxaloacetate). These flux values agree with the high protein expression levels of rTCA cycle enzymes measured in the *N. moscoviensis* proteome (Supplementary Table 2) and gene expression recently reported in the transcriptome (12). In particular, isocitrate dehydrogenase (IDH), 2-oxoglutarate:ferredoxin oxidoreductase (OFOR), and pyruvate:ferredoxin oxidoreductase (PFOR) were the most abundant TCA cycle enzymes (Supplementary Table 2). This is consistent with the large thermodynamic barriers encountered by the consecutive reactions of alpha-ketoglutarate synthase and isocitrate dehydrogenase (*Δ*G > 40 kJ/mol), as well as pyruvate:ferredoxin oxidoreductase (*Δ*G > 14 kJ/mol) (19).

Generation of reducing equivalents for carbon fixation in *N. moscoviensis* is predicted to result from reverse electron flow from the cytochrome *c* pool (3). Overall, the model predicts that approximately 3% (0.23 mmol gDW^-1^ hr^-1^) of the oxidized nitrite is used to generate quinol (largely for nitrite assimilation via octaheme nitrite reductase and reduction of fumarate via succinate dehydrogenase), 4% (0.32 mmol gDW^-1^ hr^-1^) is used to generate NADH (via complex I), and 3% (0.27 mmol gDW^-1^ hr^-1^) is used to generate reduced ferredoxin (via the 2M-type complex I) (Supplementary Figure 2). Reduction of NADP+ to NADPH, the main electron carrier for biosynthetic reactions, was predicted to occur through NAD(P)(+) transhydrogenase. While *N. moscoviensis* encodes three NAD(P) transhydrogenases, only one (NITMOv2_1092) was detected in the proteome (Supplementary Table 1). Therefore, we posit that NITMOv2_1092 is the main transhydrogenase active during chemolithoautotrophic growth.

### Confirmation of autotrophic metabolism with ^13^C-bicarbonate tracer experiments

To confirm the biosynthetic pathways predicted by genome-scale modeling, isotopic tracers combined with high-resolution metabolomics was used to follow ^13^C-labelled bicarbonate incorporation into the metabolome of *N. moscoviensis.* Cells were grown in a membrane bioreactor under steady-state conditions and ^13^C-bicarbonate was rapidly introduced into the bioreactor, which equilibrated with the dissolved inorganic carbon (DIC) pool in the liquid media to approximately 65% ^13^C enrichment. Following ^13^C-bicarbonate addition, multiple metabolite samples were collected from the bioreactor over a 2-hour time period by rapid quenching and extraction of *N. moscoviensis* cells. A time zero sample corresponding to the period immediately before ^13^C-bicarbonate addition was also collected as a control.

Consistent with the operation of the rTCA cycle, high ^13^C label incorporation was measured in phosphoenolpyruvate (PEP), acetyl-CoA, succinate, and aspartic acid (used as oxaloacetate surrogate) (Figure 5). Other TCA cycle metabolites showed considerable ^13^C enrichment, including malate, fumarate, and alpha-ketoglutarate, although to a lower extent than PEP, acetyl-CoA, succinate, and aspartic acid (Figure 5). This suggested that the ^13^C enrichment of malate, fumarate, and alpha-ketoglutarate may have been diluted by inactive pools of these metabolites that were not labeled. Further evidence for this was observed from the faster labelling of glutamate and glutamine than alpha-ketoglutarate, which was unexpected because these amino acids are derived from alpha-ketoglutarate (Figure 5; Supplementary Dataset 3). We hypothesize that these patterns may reflect substrate channelling in the rTCA cycle of *N. moscoviensis*, which has previously been observed in the TCA cycle of other organisms (20)(21)(22).

**Figure 5.**
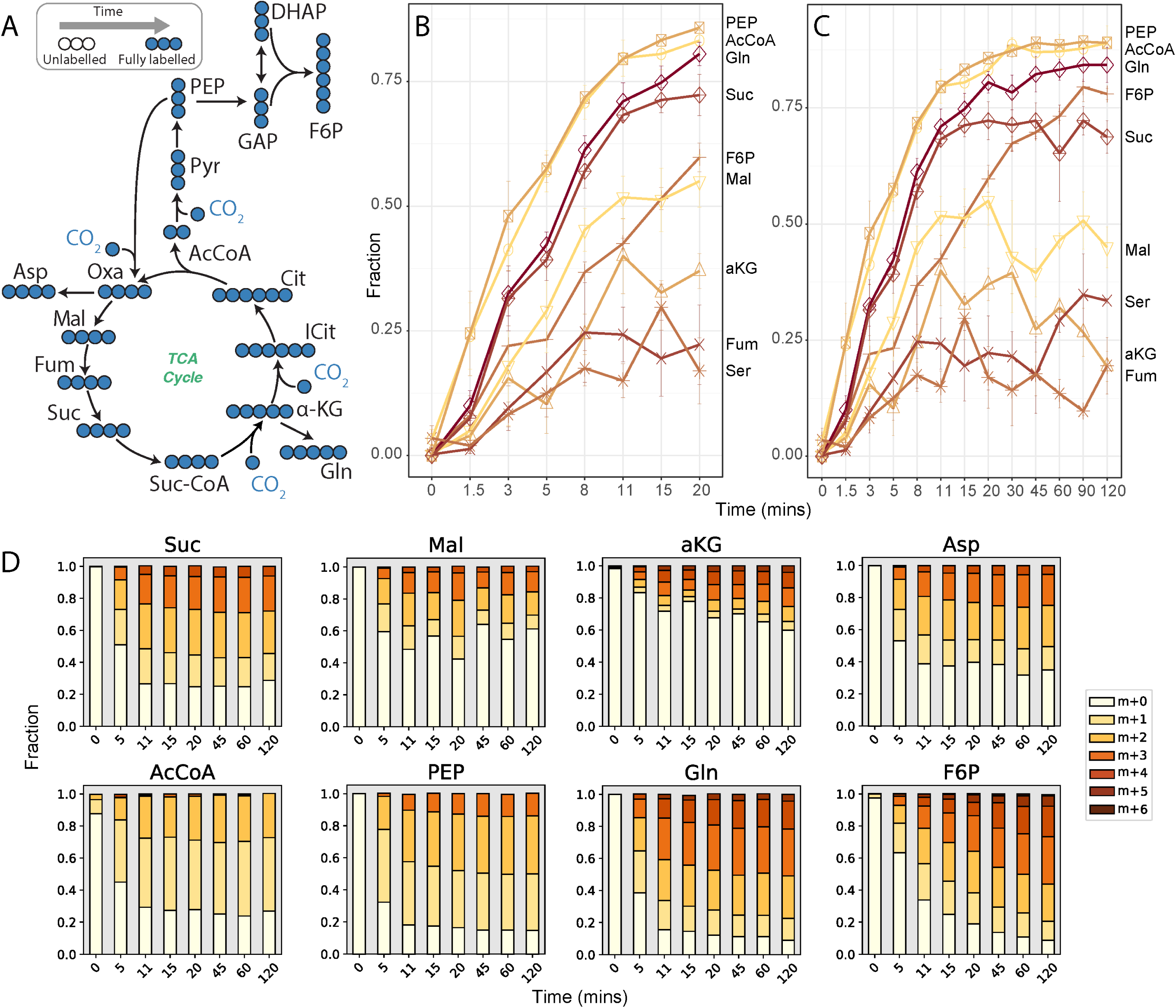
Dynamic ^13^C-labelling of selected central carbon metabolites during ^13^C-bicarbonate tracer experiments. (A) Expected ^13^C labeling of central metabolism over time. ^13^C enrichment of selected metabolites over (B) 20 minutes and (C) 120 mins. ^13^C-enrichment values were normalized to a tracer ^13^C fraction of 1. (D) Mass isotopomer distributions for selected metabolites. AcCoA: acetyl-CoA, aKG: alpha-ketoglutarate, Asp: aspartic acid, F6P: fructose 6- phosphate, Fum: fumarate, Gln: glutamine, Mal: malate, PEP: phosphoenolpyruvate, Ser: serine, Suc: succinate. All measured metabolite mass isotopomer distributions can be found in Supplementary Dataset 3.

In addition to the TCA cycle, fast labelling of 3-phosphoglycerate, fructose 6-phosphate, glucose 6-phosphate, and sedheptulose 7-phosphate was observed (Figure 5; Supplementary Dataset 3). This is consistent with the use of gluconeogenesis and the pentose phosphate pathway for synthesis of precursor metabolites in *N. moscoviensis*.

### Analysis of formatotrophic and mixotrophic growth

It has been demonstrated that *N. moscoviensis* can use formate as an energy source and that the genome contains a formate transporter (focA) and a soluble NAD^+^-reducing formate dehydrogenase (fds), which catalyses the oxidation of formate to CO_2_ with concomitant reduction of NAD^+^ to NADH (5)(23). It has also been shown that formate can also be assimilated as a carbon source (9), although the pathway for assimilation currently remains unclear. Metabolic reconstruction suggested that formate could potentially be assimilated either indirectly through oxidation to CO_2_ and reassimilation via the rTCA cycle or directly based on the presence of genes encoding a reductive glycine pathway (24) (Supplementary Dataset 1, Supplementary Table 2). Therefore, we used iNmo686 to explore the growth and metabolism of *N. moscoviensis* on formate compared to chemolithoautotrophic conditions. Growth comparisons were done on minimal NOB media with the following substrates: 1) nitrite and O_2_, 2) formate and O_2_, and 3) nitrite + formate and O_2_. For Case 2, we further examined the growth benefits of using either the reductive glycine pathway or the rTCA cycle for formate assimilation by turning on/off key reactions in the model. The formate uptake rate in the model was set equal to the nitrite uptake rate, based on previously published data (5). The model predicted that formate or formate + nitrite utilization would increase the growth rate of *N. moscoviensis* by approximately 2.6 or 3.8 fold, respectively (Figure 6). Here, electrons derived from formate oxidation were predicted to drive energy conservation via the electron transport chain, while also providing reducing equivalents for biosynthesis.

**Figure 6.**
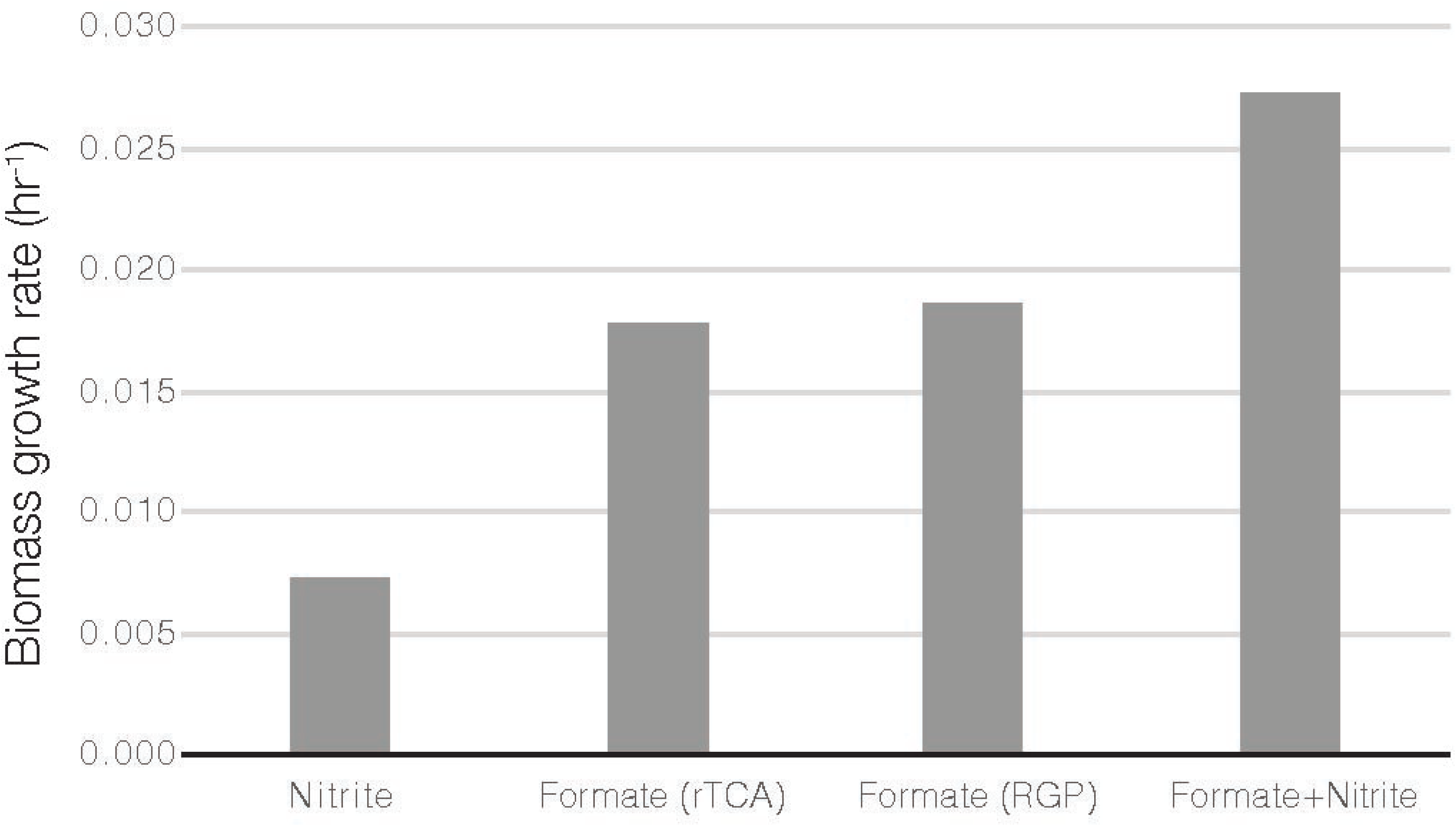
Model predictions for growth rate utilizing different substrates (nitrite and/or formate) and different formate assimilation pathways. rTCA: reductive TCA cycle, RGP: reductive glycine pathway. In all scenarios, the uptake flux of each substrate was equal to 8.5 mmol gDW^-1^ hr^-1^.

FBA also predicted that *N. moscoviensis* growth rates would improve approximately 4% by directly assimilating formate via the reductive glycine pathway versus indirectly via the rTCA cycle (Figure 6). This is because the reductive glycine pathway consumes less energy than the rTCA, as low-potential electron carriers (i.e. reduced ferredoxin) are not needed for CO_2_ fixation, and in total four electrons less are required per pyruvate molecule formed from formate instead of CO_2_ (Figure 7). In this scenario, approximately 6% of the formate would be directly assimilated via the reductive glycine pathway, rather than oxidizing all formate to CO_2_ followed by reassimilation (Supplementary Dataset 2). Congruently, all genes for the reductive glycine pathway in the genome were also found in the proteome (Supplementary Table 2) and previously published *N. moscoviensis* transcriptome (12).

**Figure 7.**
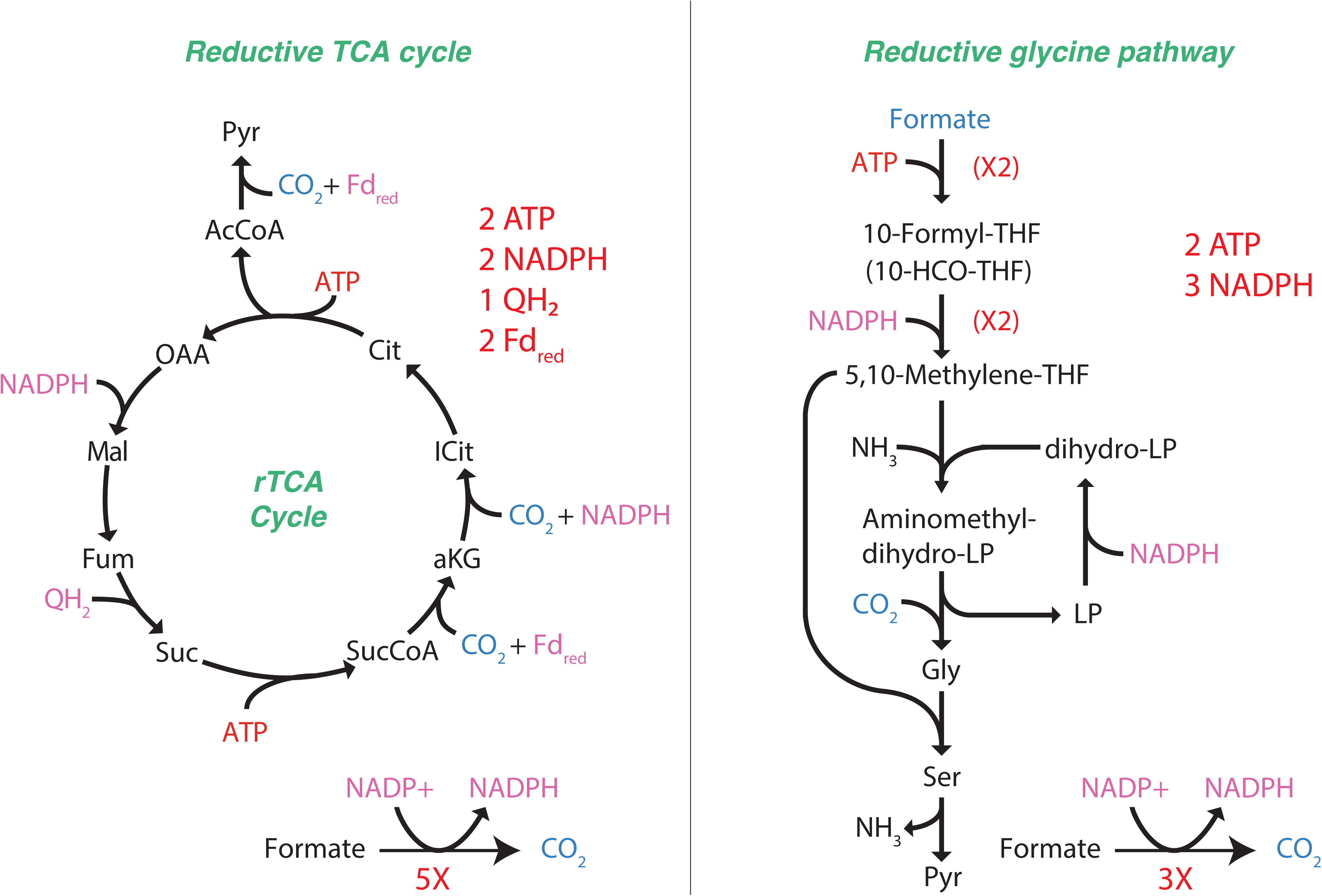
Comparison of the reductive TCA cycle with the reductive glycine pathway.

### ^13^C-formate tracer experiments suggest formate is indirectly assimilated via CO_2_ fixation

To experimentally determine the assimilation pathway of formate in *N. moscoviensis,* isotopic tracer experiments with ^13^C-formate were performed with batch cultures in sealed serum bottles. Cultures were first acclimated to growth on 1 mM unlabelled formate without the presence of nitrite for 24 hours. Subsequently 0.5 mM ^13^C-formate was added to the cultures and intracellular metabolome and gas headspace samples were collected before addition of ^13^C-formate and at 15, 30, 60, 180, and 300 mins for isotopic analysis. Consistent with ^13^C-formate oxidation to CO_2_, continuous production of ^13^C-CO_2_ in the headspace gas was observed during the experiment (Figure 8A). Intracellular formate was also measured to be ∼40% ^13^C-enriched from 15 minutes onwards, consistent with formate transport into the cell (Figure 8C). However, the majority of measured metabolites had no detectable ^13^C enrichment, expect for succinate, glutamate, glutamine, and fructose 6-phosphate (Supplementary Dataset 3, Figure 8C). Moreover, mass isotopomer distributions for these metabolites showed a consistent increase in labelled carbons overtime (Figure 8C), similar to results with ^13^C-bicarbonate experiments (Figure 3). This suggests that formate was being oxidized to CO_2_ and then assimilated via the rTCA cycle. Batch experiments with *N. moscoviensis* also observed slower growth on formate than nitrite (Figure 8B). This was tested over a range of pH conditions, as previous studies indicated that formate oxidation may have a different pH optimum than nitrite oxidation (25)(26). These results reinforce the previous findings of poor growth of *N. moscoviensis* on formate (5) and do not support the model predictions of an improved growth rate on formate.

**Figure 8.**
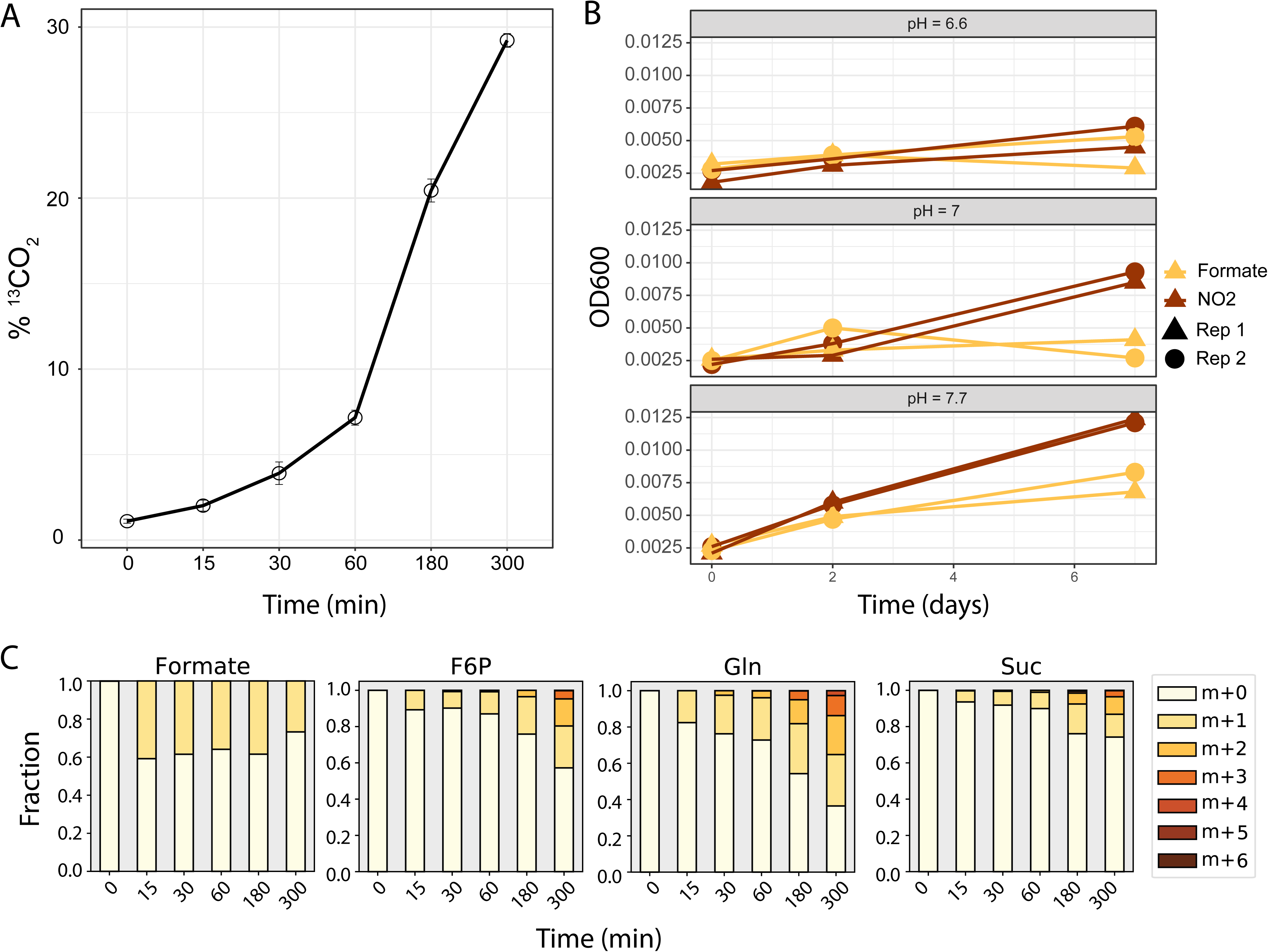
Oxidation and assimilation of ^13^C-formate by *N. moscoviensis.* (A) Production of ^13^C- CO_2_ from ^13^C-formate oxidation. (B) Growth of *N. moscoviensis* using formate (yellow) or nitrite (red) as electron donor with oxygen as the terminal electron acceptor at different pH. (C) Mass isotopomer distributions for selected metabolites during batch ^13^C-formate tracer experiments.

## Discussion

Previous studies have shown the utility of constraint-based reconstruction and analysis for exploring the metabolic capabilities of microorganisms that have an important environmental role (27)(28)(29). iNmo686 provides a framework for examining *Nitrospira* metabolism at a systems-level and will serve as a knowledge base that can be continually refined to drive understanding and improved prediction of *Nitrospira* ecophysiology. Our analysis shows that iNmo686 quantitatively predicts *N. moscoviensis* chemolithoautotrophic growth on nitrite. Flux balance analysis provided estimates for the amount of nitrite used for energy conservation and CO_2_ fixation into biomass, offering a quantitative model for linking catabolism and anabolism. The model also accurately provides quantitative estimates of carbon and electron flux distribution, reinforcing the metabolic network predicted based on the proteomic measurements from this study and from previous genomic and transcriptomic analyses of *Nitrospira* (3)(12).

^13^C metabolomic analysis revealed a complex picture for central carbon metabolism in *Nitrospira*. While our results support that *Nitrospira* uses the rTCA cycle for CO_2_ fixation under chemolithoautotrophic growth, ^13^C enrichment values revealed extensive unlabelled pools of central carbon intermediates in the metabolome (Figure 5). Because all ^13^C enrichment values increased monotonically, we reasoned that turnover of carbon storage reserves or macromolecules was not responsible for this observation. However, compartmentalization of metabolism or substrate channeling may offer a possible explanation. In cells that contain multiple pools of the same metabolite (e.g. cells with mitochondria) differences in their labelling will result in an aggregated measurement of the mixed pool because they are extracted together. If only one of those pools is actively participating in metabolism, the aggregation of both labelled and unlabelled pools will serve to dilute the overall ^13^C enrichment. While electron micrograph images suggest that no intracytoplasmic membranes or carboxosomes are present in *N. moscoviensis* cells (6)(30)(31), it is possible that separate metabolite pools may arise from certain enzymes/pathways having different subcellular locations within the cytoplasm (32). However, this requires further investigation.

An alternative mechanism to explain the low ^13^C enrichment levels of some metabolites is substrate channeling; that is, the direct passing of pathway intermediates between enzyme active sites without escaping into the cytoplasm, facilitated by noncovalent dynamic enzyme complexes (33)(34). This can result in lower than expected ^13^C enrichment values due to “leaked” metabolites that create a separate cytoplasmic pool with a much slower turnover rate (33)(34). Given that several metabolites immediately downstream of the rTCA cycle (aspartate, glutamate, glutamine) were more labelled than their precursor TCA intermediates, we suspect that rTCA cycle enzymes may interact to form a supramolecular complex, or metabolon, to efficiently transport reactants between enzyme active sites. The advantage of this is that high local substrate concentrations enable higher pathway fluxes and intermediates can be protected from the bulk phase, limiting competition between competing pathways and protecting the cell from toxicity (33)(35). Given the significant thermodynamic barriers encountered in the rTCA by the consecutive operation of alpha-ketoglutarate synthase and isocitrate dehydrogenase (*Δ*G > 40 kJ/mol), as well as pyruvate synthase (*Δ*G > 14 kJ/mol) (19), substrate channeling maybe be important for overcoming these unfavorable reactions in addition to their indirect coupling to ATP hydrolysis via succinyl-CoA synthetase and ATP citrate lyase. Further confirmation of this channeling could be obtained through *in vivo* crosslinking coupled to proteomic analysis, as has been done to investigate the malate dehydrogenase–citrate synthase–aconitase complex of the oxidative TCA cycle (21)(22).

In addition to chemolithoautotrophic growth on nitrite, genome-scale modeling allowed us to assess different hypotheses regarding formatotrophic and mixotrophic growth of *N. moscoviensis*. *Nitrospira* have been observed to oxidize formate to CO_2_ for energy conservation under laboratory conditions (5) and also to assimilate formate *in situ* during wastewater treatment (9). However, while our analysis predicted that *N. moscoviensis* should grow 2-3 times faster on formate over nitrite, batch experiments showed lower biomass yields of *N. moscoviensis* on formate, consistent with previous reports (5). This suggests that formate may be toxic to *N. moscoviensis* via an unknown mechanism. Given that different lineages of *Nitrospria* may exhibit different formate utilization efficiencies *in situ* (9), it remains to be tested whether all *Nitrospria* grow slower using formate as an electron donor or whether other environmental conditions boost formatotrophic growth.

Genome-scale modeling further allowed us to generate hypotheses on the use of formate as a carbon source for *N. moscoviensis* anabolism. While modeling predicted that small growth improvements would be achieved by directly assimilating formate via the reductive glycine pathway, *in vivo* ^13^C-formate tracer experiments demonstrated that *N. moscoviensis* adhere to their autotrophic lifestyle, reducing CO_2_ derived from formate via the rTCA cycle. This may be an evolutionary adaptation to avoid energetic costs associated with remodeling their proteome in environments where formate may only become transiently available, for example at oxic-anoxic interfaces common to the habitats of *Nitrospira* (1). Moreover, this highlights the importance of validating model predictions with experimental measurements.

In conclusion, our work provides the first genome-scale reconstruction and analysis of *Nitrospira* metabolism, offering unique insights on their versatile ecophysiology. The iNmo686 GEM provides quantitative estimates of chemolithoautotrophic growth and pathway fluxes and serves as an invaluable tool for hypothesis-driven discovery. Our ^13^C metabolomic results also provide the first insights on *Nitrospira’s in vivo* central carbon metabolism, confirming the high activity of the rTCA cycle for CO_2_ fixation. Future efforts to combine ^13^C metabolomics with genome-scale modeling should provide a valuable approach for quantitatively understanding regulation of *Nitrospira* metabolism under different environmental conditions and microbe-microbe interactions. This will further expand the systems biology framework developed in this study, ultimately leading to the systematic prediction and control of *Nitrospira* metabolism in natural and engineered ecosystems.

## Materials and Methods

### Cultivation of N. moscoviensis cells

*N. moscoviensis* M-1 was grown in NOB mineral salts medium for lithoautotrophic growth as described in Spieck and Lipski (2011) (36) with the following modifications: CaCO_3_ was replaced with CaCl_2_ ⋅ 2 H_2_O in the same concentration and the following trace element composition was used per liter of medium: 34.4 μg of MnSO_4_ ⋅ 1 H_2_O, 50 μg of H_3_BO_3_, 70 μg of ZnCl_2_, 72.6 μg of Na_2_MoO_4_ ⋅ 2 H_2_O, 20 μg of CuCl_2_ ⋅ 2 H_2_O, 24 μg of NiCl_2_ ⋅ 6 H_2_O, 80 μg of CoCl_2_ ⋅ 6 H_2_O, and 2,000 μg of FeSO_4_ ⋅ 7 H_2_O. Nitrilotriacetic acid was added equimolar to all trace elements as a complexing agent.

*N. moscoviensis* was cultivated in a 7L membrane bioreactor (MBR) inoculated with an active batch culture and operated as described in Mundinger et al., (2019) (12). The working volume of the reactor was 3 L and included pH, dissolved oxygen, temperature, and level controls (all by Applikon Biotechnology B.V., Schiedam, The Netherlands). The bioreactor was continuously sparged with Ar/CO_2_ (95%/5% v/v) and air at a rate of 10 ml/min to maintain a dissolved oxygen concentration of ∼30%. pH was controlled at 7.7 using a 1 M KHCO_3_ buffer and the reactor temperature was maintained at 39°C using a loop-type heat exchanger. The reactor was continuously stirred at 150 rpm. The reactor and all cultivation media and solutions were sterilized by autoclaving or sterile filtration prior to use and the reactor was operated aseptically to maintain culture purity. Nitrite and nitrate concentrations were check daily to ensure all nitrite was consumed stoichiometrically to nitrate (Nitrite test strips MQuant®, Merck, Darmstadt, Germany) and that nitrite was always limiting. Cultures were maintained in steady-state growth at a dilution rate of 0.006 hr^-1^ and a substrate feeding rate of 2 – 2.5 mmol NO_2_^-^ L^-1^ day^-1^.

### Genome-scale model reconstruction and analysis

The genome-scale metabolic model of *N. moscoviensis* (*iNmo686)* was reconstructed from the NCBI’s genome sequence for *N. moscoviensis* (accession number NZ_CP011801.1) using the Model SEED pipeline (37) implemented in KBase (14), followed by manual curation using the MetaCyc database (13) and available literature (1)(3)(5). The model was gap-filled manually through the addition of reactions not annotated in the genome to ensure that all biomass components could be produced on NOB minimal media. Growth and non-growth associated maintenance energy requirements were determined by plotting experimentally measured nitrite uptake rates as a function of the growth rate set during steady-state bioreactor cultivation (38). The biomass equation was derived from biomass composition measurements of *N. moscoviensis* (Supplementary Dataset 1). The model was formulated in Systems Biology Markup Language (SBML) level 3 version 1.0 and is available at GitHub (https://github.com/celawson87/Nitrospira-moscoviensis-GEM). Flux balance analysis was used to simulate *in silico* growth by solving the linear program:

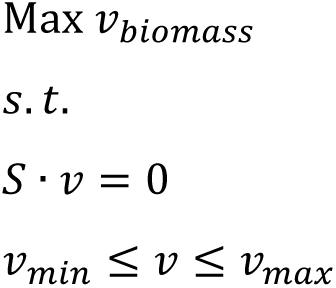

where *v_biomass_* is the flux through the biomass objective function, *S* is the stoichiometric matrix generated from the reconstruction with rows representing metabolites, columns representing reactions, and entries representing metabolite stoichiometric coefficients, *v* is the vector of steady-state reaction fluxes, and *v_min_* and *v_max_* are the minimum and maximum allowable reaction fluxes. Flux balance analysis was performed in Python version 3.7.2 using the COBRApy package (39).

The transcriptomic dataset reanalyzed here can be found under the NCBI Gene Expression Omnibus DataSet accession number GSE123406.

### Biomass composition analysis

Cultures were centrifuged (10,000 rpm, 15 mins, 4°C) to obtain cell pellets, which were subsequently freeze-dried prior to analysis. Total protein concentration was determined using the Pierce^TM^ BCA Protein Assay Kit (ThermoFisher Scientific) and amino acid composition was determined according to Carnicer et al. (2009) (40) using a Varian 920-LC high performance liquid chromatography amino acid analyzer. Total carbohydrates were determined using the phenol-sulphuric acid method (41). Total lipid content was determined via the sulfo-phospho-vanillin reaction (42) and lipid composition for *N. moscoviensis* was taken from Lipski et al. (2001) (18). Total RNA and DNA content was determined according to Benthin et al. (1991) (43). Total inorganic content was determined by combustion of freeze-dried biomass in an oven at 550°C for 12 hours. Lipid headgroup composition, ion composition, and soluble pool composition was derived from the *Escherichia coli* biomass equation reported for iAF 1260 (44).

### ^13^C isotopic tracer experiments

^13^C-labelled sodium bicarbonate was rapidly introduced (within 1 minute) into the bioreactor containing *N. moscoviensis* cells growing under steady-state conditions to a final concentration of approximately 30 mM. Following ^13^C-label introduction, samples were rapidly withdrawn from the reactor at timepoints 0, 1.5, 3, 5, 8, 11, 15, 20, 30, 45, 60, 90, and 120 minutes. Samples were immediately filtered (Millipore 0.45 µm hydrophilic nylon filter HNWPO4700) using a vacuum pump to remove extracellular media, and filters were placed face down in 1.5 ml of - 80°C extraction solvent (40:40:20 acetonitrile:methanol:water) for cell quenching and metabolite extraction. Samples were then centrifuged (10,000 rpm, 4°C, 5 mins) and 1 ml of cell-free supernatant was collected and stored at -80°C for metabolomic analysis. The time 0 min sample corresponded to the period directly before ^13^C-label addition. The ratio of ^13^C/^12^C dissolved inorganic carbon remained constant during the course of the 2-hour experimental period as determined by GC-MS analysis.

### Formate batch experiments

*N. moscoviensis* cells were harvested from a membrane bioreactor. The biomass was centrifuged 8,000 × *g*, 15 min at 25°C and washed by resuspending the cells in fresh mineral NOB media. This was repeated until no nitrite or nitrate was detectable via nitrite/nitrate test strips (Nitrite test strips MQuant®, Merck) in the culture. Subsequently, the cells were transferred to sterile 120 ml serum bottles (triplicate experiments, 1 bottle per timepoint, 6 timepoints total) containing 50 ml NOB mineral salts medium with 0.5 mM sodium formate and no nitrite but 0.187 mM NH_4_Cl as nitrogen source. Bottles were crimp sealed with a rubber stopper to allow monitoring of the gas headspace and were incubated at 39°C in the dark. Following 24 hours of acclimation, 1 mM ^13^C-sodium formate (Cambridge Isotopes Laboratories, MA, USA) was added to all incubations. At each timepoint (before addition, 15, 30, 60, 180, 300 minutes) the isotopic composition of the gas headspace was measured using GCMS and subsequently bottles corresponding to a given timepoint were sacrificed for metabolomics analysis. Bottle contents were filtered at a given timepoint using a vacuum pump and metabolites were extracted using - 80°C extraction solvent (40:40:20 acetonitrile:methanol:water) as described above.

For growth experiments, *N. moscoviensis* cells from a membrane bioreactor were harvested and washed until no nitrite or nitrate was detectable via nitrite/nitrate test strips (Nitrite test strips MQuant®, Merck) (see above). The cells were transferred to sterile 120 ml serum bottles with 50 ml NOB mineral media containing 0.187 mM NH_4_Cl as nitrogen source and a pH of 6.6, 7 or 7.7 adjusted using 1M KHCO_3_ and were sealed with a rubber stopper. The experiment was performed in duplicates (per pH value and substrate) and 5 mM sodium nitrite or 5 mM sodium formate were added to the incubations as sole energy source. Growth was monitored for 7 days using optical density measurements at wavelength 600 nm.

### Metabolomic analysis

Samples were analysed using a high-performance HPLC–MS system consisting of a Vanquish^TM^ UHPLC system (Thermo Scientific) coupled by electrospray ionization (ESI; negative polarity) to a hybrid quadrupole high-resolution mass spectrometer (Q Exactive Orbitrap, Thermo Scientific) operated in full scan mode for detection of targeted compounds based on their accurate masses. Properties of Full MS–SIM included a resolution of 140,000, AGC target of 1E6, maximum IT of 40 ms and scan range from 70 to 1,000 m/z. LC separation was achieved using an ACQUITY UPLC BEH C18 column (2.1 × 100 mm column, 1.7 μm particle size; part no. 186002352; serial no. 02623521115711, Waters). Solvent A was 97:3 water:methanol with 10 mM tributylamine (TBA) adjusted to pH 8.1–8.2 with 9 mM acetic acid. Solvent B was 100% methanol. Total run time was 25 min with the following gradient: 0 min, 5% B; 2.5 min, 5% B; 5 min, 20% B; 7.5 min, 20% B; 13 min, 55% B; 15.5 min, 95% B; 18.5 min, 95% B; 19 min, 5% B; 25 min, 5% B. Flow rate was 200 μl min^−1^. The autosampler and column temperatures were 4°C and 25°C, respectively. Mass isotopomer distributions were corrected for natural abundance using the method of Su et al., (2017)(45) and ^13^C enrichment values were calculated using the formula 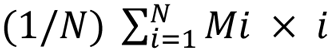, where *N* is the number of carbon atoms in the metabolite and *Mi* is the fractional abundance of the *i^th^* mass isotopomer.

### GC-MS analysis of dissolved inorganic carbon isotopic fractions

Isotopic fractions of dissolved inorganic carbon in the liquid media were measured based on a modified headspace method (46). 3 ml of liquid culture were collected from the bioreactor with a syringe and directly filtered through a 0.45 µm filter and 26G needle into a 120 ml bottle containing 1 ml 6 M HCl (strong acid) crimp sealed with a rubber stopper. Prior to adding the liquid sample, bottles and HCl were flushed with either 100% N_2_ or Ar gas to void the headspace of background CO_2_. Samples were equilibrated with the acid in the bottles for at least 1 hour at room temperature to drive all dissolved inorganic carbon into the gas phase. 50 ul gas samples were then collected in a gas tight syringe with a needle (Hamilton) from the bottle’s headspace and the isotopic fractions of ^12^CO_2_ and ^13^CO_2_ were determined using a gas chromatograph (Agilent 6890 equipped with 6 ft Porapak Q and molecular sieve columns) coupled with a mass spectrometer (GC-MS) (Agilent 5975C GC MSD; Agilent, Santa Clara, CA).

### Large-scale discovery proteomics of whole cell and membrane fraction - whole cell fraction

#### Sample preparation and proteolytic digestion

Biomass was harvested by centrifugation, washed once with Tris-EDTA buffer, pH 7.7, and frozen using liquid N_2_. Cell pellets were stored at -80°C until further processing. For protein extraction, cell pellets (445 mg wet weight) were resuspended in 5 ml 50 mM triethylammonium bicarbonate (TEAB) buffer, 1% sodium deoxycholate, pH 8, and mixed 1:1 with B-PER reagent (Thermo Fisher Scientific, Waltham, MA, USA). Cells were lysed by two rounds of sonification (Sonifier B-12 with microtip, Branson Sonic Power Company, Danbury, CT, USA) at setting 6 for 30 sec, intermitted by cooling on ice. Cell debris and unopened cells were separated from the sample by centrifugation (14,000 x g, RT, 10 min). Proteins were precipitated by addition of 250 µl 100% (w/v) trichloroacetic acid (TCA) to 1 ml of sample and incubation for 20 min on ice. Proteins were collected by centrifugation (14,000 rpm, RT, 5 min) and washed two times with ice cold acetone. Next, the protein pellet was resuspended in 400 µl 50 mM TEAB buffer aided by a heating step at 40°C, 1h, with intermitted vortexing. 100 µl of the protein extract were diluted 5 times with 6 M urea, 200 mM ammonium bicarbonate (ABC) buffer followed by vortexing and a heating step for 20 min at 45°C, 1450 rpm and 25 min of sonification. Acetonitrile (ACN) was added to a final concentration of 50%, followed by another heating step. A 200 µl aliquot of the sample was reduced by the addition of dithiothreitol (DTT) to a final concentration of 2.3 mM followed by incubation at 37°C, 1h. Next, the sample was alkylated by the addition of iodoacetamide (IAM) to a final concentration of 3.75 mM followed by incubation in the dark at room temperature for 30 min. The sample was diluted with 200mM ABC buffer to a concentration of urea below 1 M. Lastly, an 150 µL aliquot of the protein sample was digested with 5 µg of sequencing grade trypsin (Promega, Madison, WI, USA) overnight at 37°C.

### Large-scale discovery proteomics of whole cell and membrane fraction - membrane fraction

#### Sample preparation and proteolytic digestion

Biomass was harvested by centrifugation, washed once with TE buffer and frozen using liquid N_2_. Cell pellets were stored at -80°C until further processing. For protein extraction, cell pellets (1.95 g wet weight) were resuspended in 13 ml 50 mM TEAB buffer, 1% Sodium Deoxycholate, pH 8, and mixed 1:1 with B-PER reagent (Thermo Fisher Scientific). Cells were lysed by three passages through a French press at 138 MPa pressure. Cell debris and unopened cells were pelleted from the sample by centrifugation (5,000 x g, 4°C, 15 min). The membrane fraction was prepared using ultracentrifugation (45,000 rpm, 4 °C, 1h; Optima-90 ultracentrifuge, Beckman-Coulter, Brea, CA, USA) and washed once in 200 mM ABC buffer, followed by another ultracentrifugation step to collect the membrane pellet. Finally, the membrane pellet was resuspended in 400 µl 200 mM ABC, 1% DDM and membrane proteins solubilized overnight in an orbital shaker at 4 °C. The sample was clarified by centrifugation (20,000 x g, RT, 15 min). Urea was added to a final concentration of 6 M into the sample. The sample was reduced by the addition of DTT to a final concentration of 2.3 mM followed by incubation at 37°C for 1h. Next, the sample was alkylated by the addition of IAM to a final concentration of 3.75 mM, followed by incubation in the dark at RT, 30 min. The sample was diluted to a concentration of 1 mg/ml with 200 mM ABC, 1% DDM, 6 M Urea. Five aliquots of 100 µg protein each were digested or double digested using 2 µg of each selected proteinase, all digestion steps were performed at 37°C. The following different proteinases and combinations were tested: (1) LysC (Pierce), 4 h; (2) Trypsin (Promega), 4 h; (3) Chymotrypsin (Pierce), overnight; (4) LysC, 4h followed by Trypsin, overnight (5) Trypsin, 4h followed by Chymotrypsin, overnight. In digests with Trypsin and Chymotrypsin, ACN was added to a final concentration of 20% and the concentration of urea in the sample was diluted to ≤ 1 M by the addition of ABC buffer.

#### Solid phase extraction for membrane protein digest

Proteolytic digests were desalted using an Oasis HLB 96 well plate (Waters, Milford, MA, USA) according to the manufacturer’s protocol. The purified peptide eluate was further dried using a speed-vacuum concentrator.

#### Large-scale shot-gun proteomics

All dried peptide fractions were resuspended in H_2_O containing 3% acetonitrile and 0.1% formic acid using mild vortexing. An aliquot of every sample corresponding to approx. 100-200 ng protein digest was analysed in duplicates using an one dimensional shot-gun proteomics approach (47). Briefly, 1 µL of sample was injected to a nano-liquid-chromatography system consisting of an ESAY nano LC 1200, equipped with an Acclaim PepMap RSLC RP C18 separation column (50 µm x 150 mm, 2µm and 100A), and an QE plus Orbitrap mass spectrometer (Thermo). The flow rate was maintained at 300 nL/min over a linear gradient using H_2_O containing 0.1% formic acid as solvent A, 80% acetonitrile in H_2_O and 0.1% formic acid as solvent B. The soluble protein extract fractions were analysed using a gradient from 5% to 30% solvent B over 90 minutes, and finally to 75% B over 25 minutes. The soluble membrane protein fractions were analysed using a shorter gradient from 4 to 30% B over 32.5 minutes followed by a second step to 65% B over 12.5 minutes, and data were acquired in total over 50 minutes. In either case, the Orbitrap was operated in data depended acquisition mode acquiring peptide signals form 350 - 1400 m/z at 70K resolution, where the top 10 signals were isolated at a window of 2.0 m/z and fragmented using a NCE of 30. The AGC target was set to 1e5, at a max IT of 54 ms and 17.5K resolution.

#### Database search and data processing

Raw shot-gun proteomics data from membrane proteins and soluble protein extract fractions were analysed and combined using PEAKS Studio 8.5 (Bioinformatics Solutions Inc., http://www.bioinfor.com). Database search was performed allowing 20 ppm parent ion and 0.02 Da fragment mass error tolerance. Search conditions further considered 3 missed cleavages for the respective enzymes used, carbamidomethylation as fixed and methionine oxidation and N/Q deamidation as variable modifications. Peptide spectra were matched against a *N. moscoviensis* sequence database (Uniprot TrEMBL, June 2018, Tax ID 42253), which was modified by filtering out duplicated entries. Database search included the GPM crap contaminant database (https://www.thegpm.org/crap/) and a decoy fusion for determining false discovery rates. Peptide spectrum matches were filtered against 1% false discovery rate (FDR) and protein identifications with 2 or more unique peptides were considered as significant hit. For the prefractionated whole cell proteome, proteins were counted if they were significantly identified in at least one of the subfractions.

### Transmembrane helix predictions

The combined transmembrane topology and signal peptide predictor tool Phobius (http://phobius.sbc.su.se, (48)) was employed for the prediction of membrane proteins. Since this tool does not take into account transmembrane helices that overlap with signal peptides, all proteins with one or more TMH prediction we defined as membrane proteins.

### Transcriptome analysis

In order to relate the proteome data to the previously published transcriptome of *N. moscoviensis* under stable cultivation conditions grown on nitrite (12), we reanalyzed the transcriptome data (GEO Series accession number GSE123406) using the *N. moscoviensis* genome annotation version CP011801.1 as reference, from which the Uniprot protein sequences were derived (5). The transcriptomic reanalysis was performed as in Mundinger et al. 2019 (12), in short quality filtered raw reads from IonTorrent PGM sequencing (minimum quality score of 0.05, maximum sequencing length of 300 bp, allowing two ambiguous nucleotides) were mapped to the *N. moscoviensis* genome (NCBI accession number CP011801.1) using the mapping tool BBMap v35.92 developed by Bushnell B., (https://sourceforge.net/projects/bbmap/) and counted using featureCounts (49) with the parameters ‘minid-0.95’ with ‘fracOverlap-0.9’ for a minimum alignment identity of 95% over 90% of the read length and ‘ambig=random’ to assign reads with multiple top-scoring mapping locations randomly to a single location.

Since the genome includes several duplicated genes, the 4790 protein coding CDS correspond to only 4733 uniprot entries for non-identical proteins. To directly compare transcription to proteome data, the transcriptome reads of those identical genes were thus summed up.

Gene expression levels of CDS were compared by ranking them from high to low based on their reads per kilobase per million reads (RPKM) values. Additionally the log_2_-fold to median was calculated for genome-wide visualizations of the gene expression levels in a circos plot (50).

## Acknowledgements

The authors would like to acknowledge Laura Hesp and Maartje van Kessel for assistance with the ^13^C tracer experiments and Joshua Hamilton for helpful feedback on genome-scale model reconstruction. Funding was provided by the National Science Foundation (CBET-1435661, CBET-1803055 and MCB-1518130), the Netherlands Organization for Scientific Research (Grants 016.Vidi.189.050, VI.Veni.192.086 and SIAM Gravitation Grant 024.002.002), the European Research Council (ERC Advanced Grant Ecomom 339880), a Wisconsin Distinguished Graduate Fellowship, a Postgraduate Scholarship-Doctoral (PGS-D) by the National Sciences and Engineering Research Council of Canada (NSERC), and the UW-Madison Office of the Vice Chancellor for Research and Graduate Education through the Microbiome Initiative. This research was performed in part using the Wisconsin Energy Institute computing cluster, which is supported by the Great Lakes Bioenergy Research Center as part of the U.S. Department of Energy Office of Science (DE-SC0018409).

## Contributions

C.E.L., S.L., K.D.M., and D.R.N. designed the study. C.E.L. built the models and C.E.L. and T.B.J. performed the metabolomic analysis. C.E.L., A.B.M., and H.K. performed the ^13^C isotopic tracer experiments. A.B.M., M.P., and H.K. performed the proteomic analysis. H.K. performed the ^13^C-formate batch experiments. C.E.L. and C.W. performed the biomass compositional analysis. C.E.L. wrote the manuscript. All authors provided valuable feedback and edits on the manuscript.

## Conflict of interest statement

The authors declare no conflicts of interest.

## Reference

1. Daims H, Lücker S, Wagner M. 2016. A New Perspective on Microbes Formerly Known as Nitrite-Oxidizing Bacteria. Trends Microbiol 24:699–712.

2. Daims H, Nielsen JL, Nielsen PH, Schleifer K-H, Wagner M. 2001. In Situ Characterization of Nitrospira-Like Nitrite-Oxidizing Bacteria Active in Wastewater Treatment Plants. Appl Environ Microbiol 67:5273 LP – 5284.

3. Lücker S, Wagner M, Maixner F, Pelletier E, Koch H, Vacherie B, Rattei T, Damsté JSS, Spieck E, Le Paslier D, Daims H. 2010. A Nitrospira metagenome illuminates the physiology and evolution of globally important nitrite-oxidizing bacteria. Proc Natl Acad Sci U S A 107:13479–84.

4. Koch H, Galushko A, Albertsen M, Schintlmeister A, Spieck E, Richter A, Nielsen PH, Wagner M, Daims H, Gruber-Dorninger C, Lücker S, Pelletier E, Le Paslier D, Spieck E, Richter A, Nielsen PH, Wagner M, Daims H. 2014. Growth of nitrite-oxidizing bacteria by aerobic hydrogen oxidation. Science 345:1052–4.

5. Koch H, Lücker S, Albertsen M, Kitzinger K, Herbold C, Spieck E, Nielsen PH, Wagner M, Daims H. 2015. Expanded metabolic versatility of ubiquitous nitrite-oxidizing bacteria from the genus Nitrospira. Proc Natl Acad Sci 112:201506533.

6. Ehrich S, Behrens D, Lebedeva E V, Ludwig W, Bock E. 1995. A New Obligately Chemolithoautotrophic, Nitrite-Oxidizing Bacterium, Nitrospira-Moscoviensis Sp-Nov and Its Phylogenetic Relationship. Arch Microbiol 164:16–23.

7. van Kessel MAHJ, Speth DR, Albertsen M, Nielsen PH, Op den Camp HJM, Kartal B, Jetten MSM, Lücker S. 2015. Complete nitrification by a single microorganism. Nature 528:555–559.

8. Daims H, Lebedeva EV, Pjevac P, Han P, Herbold C, Albertsen M, Jehmlich N, Palatinszky M, Vierheilig J, Bulaev A, Kirkegaard RH, von Bergen M, Rattei T, Bendinger B, Nielsen PH, Wagner M. 2015. Complete nitrification by Nitrospira bacteria. Nature 528:504–509.

9. Gruber-Dorninger C, Pester M, Kitzinger K, Savio DF, Loy A, Rattei T, Wagner M, Daims H. 2014. Functionally relevant diversity of closely related Nitrospira in activated sludge. ISME J 9:643–55.

10. Orth JD, Thiele I, Palsson BØ. 2010. What is flux balance analysis? Nat Biotechnol 28:245–8.

11. Jang C, Chen L, Rabinowitz JD. 2018. Metabolomics and Isotope Tracing. Cell 173:822– 837.

12. Mundinger AB, Lawson CE, Jetten MSM, Koch H, Lücker S. 2019. Cultivation and Transcriptional Analysis of a Canonical Nitrospira Under Stable Growth Conditions 10:1– 15.

13. Caspi R, Billington R, Fulcher CA, Keseler IM, Kothari A, Krummenacker M, Latendresse M, Midford PE, Ong Q, Ong WK, Paley S, Subhraveti P, Karp PD. 2017. The MetaCyc database of metabolic pathways and enzymes. Nucleic Acids Res 46:D633– D639.

14. Arkin AP, Cottingham RW, Henry CS, Harris NL, Stevens RL, Maslov S, Dehal P, Ware D, Perez F, Canon S, Sneddon MW, Henderson ML, Riehl WJ, Murphy-Olson D, Chan SY, Kamimura RT, Kumari S, Drake MM, Brettin TS, Glass EM, Chivian D, Gunter D, Weston DJ, Allen BH, Baumohl J, Best AA, Bowen B, Brenner SE, Bun CC, Chandonia J-M, Chia J-M, Colasanti R, Conrad N, Davis JJ, Davison BH, DeJongh M, Devoid S, Dietrich E, Dubchak I, Edirisinghe JN, Fang G, Faria JP, Frybarger PM, Gerlach W, Gerstein M, Greiner A, Gurtowski J, Haun HL, He F, Jain R, Joachimiak MP, Keegan KP, Kondo S, Kumar V, Land ML, Meyer F, Mills M, Novichkov PS, Oh T, Olsen GJ, Olson R, Parrello B, Pasternak S, Pearson E, Poon SS, Price GA, Ramakrishnan S, Ranjan P, Ronald PC, Schatz MC, Seaver SMD, Shukla M, Sutormin RA, Syed MH, Thomason J, Tintle NL, Wang D, Xia F, Yoo H, Yoo S, Yu D. 2018. KBase: The United States Department of Energy Systems Biology Knowledgebase. Nat Biotechnol 36:566–569.

15. Bar-even A, Noor E, Flamholz A, Milo R. 2013. Design and analysis of metabolic pathways supporting formatotrophic growth for electricity-dependent cultivation of microbes. Biochim Biophys Acta 1827:1039–1047.

16. Chadwick GL, Hemp J, Fischer WW, Orphan VJ. 2018. Convergent evolution of unusual complex I homologs with increased proton pumping capacity : energetic and ecological implications. ISME J 12:2668–2680.

17. Feist AM, Palsson BO. 2010. The biomass objective function. Curr Opin Microbiol 13:344–349.

18. Lipski A, Spieck E, Makolla A, Altendorf K. 2001. Fatty Acid Profiles of Nitrite-oxidizing Bacteria Reflect theirPhylogenetic Heterogeneity. Syst Appl Microbiol 24:377– 384.

19. Bar-even A, Flamholz A, Noor E, Milo R. 2012. Thermodynamic constraints shape the structure of carbon fi xation pathways. Biochim Biophys Acta 1817:1646–1659.

20. Sumegi B, Sherry AD, Malloy CR. 1990. Channeling of TCA cycle intermediates in cultured Saccharomyces cerevisiae. Biochemistry 29:9106–9110.

21. Wu F, Minteer S. 2015. Krebs Cycle Metabolon : Structural Evidence of Substrate Channeling Revealed by Cross-Linking and Mass Spectrometry **. Angew Chem Int Ed 54:1851–1854.

22. Zhang Y, Beard KFM, Swart C, Bergmann S, Krahnert I, Nikoloski Z, Graf A, Ratcliffe RG, Sweetlove LJ, Fernie AR, Obata T. 2017. Protein-protein interactions and metabolite channelling in the plant tricarboxylic acid cycle. Nat Commun 8:15212.

23. Oh J, Bowien B. 1998. Structural Analysis of the fds Operon Encoding the NAD+-linked Formate Dehydrogenase of Ralstonia eutropha. J Biol Chem 273:26349–26360.

24. Sánchez-Andrea I, Guedes IA, Hornung B, Boeren S, Lawson CE, Sousa DZ, Bar-Even A, Claassens NJ, Stams AJM. 2020. The reductive glycine pathway allows autotrophic growth of Desulfovibrio desulfuricans. Nat Commun 11:5090.

25. Van Gool A, Laudelout H. 1966. Formate utilization by Nitrobacter wibogradskyi. Biochim Biophys Acta - Gen Subj 127:295–301.

26. O’Kelley JC, Nason A. 1970. Particulate formate oxidase from Nitrobacter agilis. Biochim Biophys Acta - Bioenerg 205:426–436.

27. Stolyar S, Van Dien S, Hillesland KL, Pinel N, Lie TJ, Leigh J A, Stahl D A. 2007. Metabolic modeling of a mutualistic microbial community. Mol Syst Biol 3:92.

28. Zhuang K, Izallalen M, Mouser P, Richter H, Risso C, Mahadevan R, Lovley DR. 2011. Genome-scale dynamic modeling of the competition between Rhodoferax and Geobacter in anoxic subsurface environments. ISME J 5:305–316.

29. Mellbye BL, Giguere AT, Murthy GS, Bottomley PJ, Sayavedra-Soto LA, Chaplen FWR. 2018. Genome-Scale, Constraint-Based Modeling of Nitrogen Oxide Fluxes during Coculture of Nitrosomonas europaea and Nitrobacter winogradskyi. mSystems 3:e00170–17.

30. Off S, Alawi M, Spieck E. 2010. Enrichment and Physiological Characterization of a Novel Nitrospira-Like Bacterium Obtained from a Marine Sponge. Appl Environ Microbiol 76:4640 LP – 4646.

31. Nowka B, Daims H, Spieck E. 2015. Comparison of oxidation kinetics of nitrite-oxidizing bacteria: Nitrite availability as a key factor in niche differentiation. Appl Environ Microbiol 81:745–753.

32. Meyer P, Cecchi G, Stolovitzky G. 2014. Spatial localization of the first and last enzymes effectively connects active metabolic pathways in bacteria. BMC Syst Biol 8:131.

33. W FP, Williams TCR, Sweetlove LJ, Ratcliffe RG. 2011. Capturing Metabolite Channeling in Metabolic. Plant Physiol 157:981–984.

34. Sweetlove LJ, Fernie AR. 2018. substrate channelling in metabolic regulation. Nat Commun 9:2136.

35. Bulutoglu B, Garcia KE, Wu F, Minteer SD, Banta S. 2016. Direct Evidence for Metabolon Formation and Substrate Channeling in Recombinant TCA Cycle Enzymes. ACS Chem Biol 11:2847–2853.

36. Spieck E, Lipski A. 2011. Cultivation, Growth Physiology, and Chemotaxonomy of Nitrite-Oxidizing Bacteria, p. 109–130. In Klotz, MGBT-M in E (ed.), Research on Nitrification and Related Processes, Part A. Academic Press.

37. Henry CS, DeJongh M, Best AA, Frybarger PM, Linsay B, Stevens RL. 2010. High- throughput generation, optimization and analysis of genome-scale metabolic models. Nat Biotechnol 28:977–82.

38. Varma A, Palsson BO. 1994. Stoichiometric flux balance models quantitatively predict growth and metabolic by-product secretion in wild-type Escherichia coli W3110. Appl Environ Microbiol 60:3724–3731.

39. Ebrahim A, Lerman JA, Palsson BØ, Hyduke DR. 2013. COBRApy: COnstraints-Based Reconstruction and Analysis for Python. BMC Syst Biol 7:74.

40. Carnicer M, Baumann K, Töplitz I, Sánchez-Ferrando F, Mattanovich D, Ferrer P, Albiol J. 2009. Macromolecular and elemental composition analysis and extracellular metabolite balances of Pichia pastoris growing at different oxygen levels. Microb Cell Fact 8:65.

41. Herbert D, Phipps PJ, Strange RE. 1971. Chemical Analysis of Microbial Cells, p. 209– 344. In Norris, JR, Ribbons, DWBT-M in M (eds.), Methods in Microbiology. Academic Press.

42. Izard J, Limberger RJ. 2003. Rapid screening method for quantitation of bacterial cell lipids from whole cells. J Microbiol Methods 55:411–418.

43. Benthin S, Nielsen J, Villadsen J. 1991. A Simple and reliable method for the determination of cellular RNA content. Biotechnol Tech 5:39–42.

44. Feist AM, Henry CS, Reed JL, Krummenacker M, Joyce AR, Karp PD, Broadbelt LJ, Hatzimanikatis V, Palsson BØ. 2007. A genome-scale metabolic reconstruction for Escherichia coli K-12 MG1655 that accounts for 1260 ORFs and thermodynamic information. Mol Syst Biol 3:121.

45. Su X, Lu W, Rabinowitz JD. 2017. Metabolite Spectral Accuracy on Orbitraps. Anal Chem 89:5940–5948.

46. Åberg J, Wallin B. 2014. Evaluating a fast headspace method for measuring DIC and subsequent calculation of pCO2 in freshwater systems. Inl Waters 4:157–166.

47. Köcher T, Pichler P, Swart R, Mechtler K. 2012. Analysis of protein mixtures from whole-cell extracts by single-run nanoLC-MS/MS using ultralong gradients. Nat Protoc 7:882.

48. Käll L, Krogh A, Sonnhammer ELL. 2004. A combined transmembrane topology and signal peptide prediction method. J Mol Biol 338:1027–1036.

49. Liao Y, Smyth GK, Shi W. 2014. FeatureCounts: An efficient general purpose program for assigning sequence reads to genomic features. Bioinformatics 30:923–930.

50. Krzywinski M, et al, Schein J, Birol I, Connors J, Gascoyne R, Horsman D, Jones SJ, Marra MA. 2009. Circos: An information aesthetic for comparative genomics. Genome Res 19:1639–1645.

